# Locally-invasive, castrate-resistant prostate cancer in a Pten/Trp53 double knockout mouse model of prostate cancer monitored with non-invasive bioluminescent imaging

**DOI:** 10.1101/2020.04.23.057661

**Authors:** Courtney Yong, Devon L. Moose, Nadine Bannick, Marion Vanneste, Robert Svensson, Patrick Breheny, James A. Brown, Michael B. Cohen, Michael D. Henry

## Abstract

Here we have improved an existing mouse model of prostate cancer based on prostate-specific deletion of *Pten* and *Trp53* by incorporating a Cre-activatable luciferase reporter. By coupling the deletion of those genes to the activation of a luciferase reporter, we were able to monitor tumor burden non-invasively over time. We show that, consistent with previous reports, deletion of both *Pten* and *Trp53* on a C57BL/6 background accelerates tumor growth and results in both the loss of androgen receptor expression and castrate resistant tumors as compared with loss of *Pten* alone. Loss of *Trp53* results in the development of sarcomatoid histology and the expression of markers of epithelial-to-mesenchymal transition Zeb1 and vimentin, with kinetics and penetrance dependent on whether one or both alleles of *Trp53* were deleted. Homozygous deletion of *Trp53* and *Pten* resulted in uniformly lethal disease by 25 weeks. While we were able to detect locally invasive disease in the peritoneal cavity in aggressive tumors from the double knockout mice, we were unable to detect lymphatic or hematogenous metastatic disease in lymph nodes or at distant sites.

## Introduction

Prostate cancer is the most commonly diagnosed malignancy in men worldwide (1). While the 5-year survival rate is high overall and has improved with prostate specific antigen (PSA) screening guidelines and new treatment modalities, it is still the second leading cause of cancer deaths for men in the United States (2). Although most prostate cancers are found while the disease is still localized, for those with metastatic dissemination the 5-year survival drops to approximately 30% (2). Two of the most commonly altered pathways in metastatic prostate cancer are PI3K-AKT and p53. The PI3K-AKT axis is hyperactive in prostate cancer through the deletion of its negative regulator and tumor suppressor gene, phosphatase and tensin homolog on chromosome 10 (*PTEN*). *PTEN* mutations occur at a frequency of approximately 17% in primary tumors and is increased to near 40% in metastatic tumors (3, 4). Mutations in and amplification of the PI3K catalytic subunits are also observed. *TP53* mutations occur in approximately 7% of primary tumors and 50% of metastases (3, 4). Other mutations that occur at an elevated frequency in metastatic disease are in the retinoblastoma, *MYC*, and androgen receptor pathways (3, 4).

The first genetically-engineered mouse model for prostate cancer was the TRAMP model (5). TRAMP mice are engineered for prostate-specific expression of both the large and small T-antigen, which results in the inhibition of *Trp53* and *Rb1* as well as the activation of AKT. While this model exhibits metastasis, its pathology more closely reflects rare high-grade, neuroendocrine-like disease, i.e. small cell carcinoma (6, 7). In an attempt to build better models of prostate cancer, investigators have utilized prostate-specific and conditional gene knockout strategies (8).

Homozygous deletion of *Pten* using Cre recombinase controlled by a prostate-specific probasin promoter initially demonstrated lethal metastatic prostate cancer in mice (5). However, subsequent efforts found indolent disease in prostate-specific *Pten* knockout mice extensively backcrossed onto the C57BL/6 background (9, 10). Chen, *et al*. showed that combining deletion of *Trp53* with *Pten* on a C57BL/6 background unleashed a more aggressive, lethal phenotype with mice succumbing to bulky, locally invasive tumors (10). Martin, *et al*. expanded on those findings and demonstrated that the prostate specific dual deletion of *Trp53* and *Pten* resulted in rapid progression from prostate intra-epithelial neoplasia to adenocarcinoma and ultimately sarcomatoid pathology, coupled with a progressive decrease in androgen receptor staining (11). More recently, the Goodrich group has combined deletion of *Rb1* with *Pten* and/or *p53* resulting in aggressive, metastatic disease including bone metastasis (12). The addition of a *Trp53* deletion conferred castrate-resistance to the *Rb1; Pten* knockout tumors (12). However, with the deletion of *Rb1* in this model the resulting disease more closely resembled high-grade neuroendocrine prostate cancer.

We have previously described a prostate-specific *Pten* knockout mouse with a genetically engineered luciferase reporter that allowed quantitative, noninvasive monitoring of disease progression (9). However, as mentioned above, this model did not develop metastatic disease, and castration resulted tumor regression with no relapse. Here we have combined loss of *Trp53* with *Pten* with a luciferase reporter on an extensively backcrossed albino C57BL/6 background to facilitate bioluminescence imaging (BLI). We show that these mice rapidly develop fatal, bulky, locally invasive tumors. We also note the consistent development of sarcomatoid pathology and epithelial-to-mesenchymal transition (EMT)-like features in these tumors. Despite the presence of these features, we did not detect disseminated metastatic disease via BLI or histologic analysis, supporting previous findings with this model.

## Materials and Methods

### Mouse strains and genotyping

All animal procedures were performed under approval from the Institutional Animal Care and Use Committee at the University of Iowa, AP #1302028. Animals were housed in standard conditions following guidelines outlined in PHS Animal Welfare Assurance (D16-00009, A3021-01) and United States Department of Agriculture (USDA No. 42-R-0004). University of Iowa facilities have been accredited by AAALACi (#000833) since November 1994. We crossed homozygous albino C57BL/6J-Tyr^c-2J^/J ROSA26 LSL-Luc Pten^fl/fl^ mice with heterozygous B6.129P2-Trp53^tm1Brn^/J, as well as with B6.Cg-Tg(Pbsn-cre)4Prb (Pb-Cre4^+^) mice obtained from the NIH Mouse Models of Human Cancer Consortium. The mice were selected for Pten^fl/fl^ and either Trp53^fl/+^ orTrp53^fl/fl^. Once the desired alleles were obtained, mice were backcrossed to C57BL/6J-Tyr^c-2J^ mice for six generations, selecting for albino-coat color offspring. Mice were genotyped for Cre, floxed alleles of Pten, Trp53, and luciferase as described previously (13-16). Three groups of males were generated yielding 15 wild type Trp53 mice with homozygous Pten deletion (WT), 17 mice heterozygous for Trp53 with homozygous Pten deletion (HET), and 16 double knockout mice (DKO) for a total of 48 mice. Additionally, 5 more DKO mice had undergone surgical castration to study the response to androgen deprivation. Mice were examined and imaged starting at five weeks of age. Mouse health was monitored 3 times weekly and mice were euthanized by gas CO2 followed by cervical dislocation if body scores were 2 or less (17), or if they displayed signs of declining health such as decreased mobility.

### Bioluminescence imaging

All bioluminescence imaging (BLI) was performed using an Ami X imager from Spectral Instruments Imaging (Tuscon, AZ). Animals were anesthetized in a chamber with 3.5% isoflurane, and then 150 mg/kg D-luciferin (VivoGlo, Promega, Madison, WI) was injected intraperitoneally. Mice were then placed into the imager while maintaining anesthesia and an image was obtained five minutes after D-luciferin injection. Exposure time was five minutes per image. AMIView Imaging Software, version 1.5.0 was used to analyze the region of interest and measure photon flux (photons/sec/cm^2^/sr). Images were obtained on *ex vivo* necropsy tissues shortly after necropsy with an additional five-minute exposure time.

### Mouse examinations

All mice were examined and weighed bi-weekly. Examination included palpation of the abdomen and pelvis and evaluation of overall health and hygiene. Mice were imaged bi-weekly. Mice were euthanized according to pre-determined endpoints, including 20% loss of starting body weight, decreased mobility, lack of grooming, or other morbidity. Homozygous *Pten* deleted, wild type *Trp53* mice were euthanized alongside double knockout mice regardless of health status as an age-matched control. Additional cohort of *Pten* knockout mice were aged and euthanized alongside *Trp53* heterozygous mice as age matched controls for those mice. Necropsy included removal of the primary prostate tumors, imaging of the remainder of the animal, and subsequent removal of any bioluminescent tissues. Tissues were fixed in 10% neutral buffered formalin at 4°C for 24-48 hours and then dehydrated and embedded in paraffin.

### Histopathology

Formalin-fixed, paraffin-embedded tissues were sectioned and stained using standard hematoxylin and eosin stains. Immunohistochemistry was performed on serial sections with the following antibodies: cytokeratin pan antibody cocktail (ThermoFisher, Waltham, MA; catalog #MA5-13203), androgen receptor (AR) (AbCam, Cambridge, UK; catalog #ab133273), vimentin (Cell Signaling Technology, Danver, MA; catalog #5741S), and Zeb1 (obtained as a generous gift from the Richer lab at the University of Colorado) (18). Immunohistochemistry protocol included a heat-based antigen retrieval in 0.1 mM citrate buffer (pH 6.0), quench with 10% hydrogen peroxide, and blocking with 5% goat serum for all antibodies except for AR, which was blocked with 1.5% horse serum. The primary antibodies were then introduced at dilutions of 1:100 for cytokeratin, 1:250 for AR, 1:100 for vimentin, and 1.5:1000 for Zeb1. Slides were finally incubated in either horseradish peroxidase-conjugated (HRP) goat anti-rabbit (Jackson ImmunoResearch Laboratories, West Grove, PA; catalog #AB_2307391) or EnVision+ HRP anti-mouse (Dako, Carpinteria, CA; catalog #K400111-2) secondary antibodies and developed using DAKO DAB. The slides were interpreted by a single pathologist with expertise in rodent models of prostate cancer who was blinded to genotype. Analysis for EMT status was determined by overlapping regions of Zeb1 and Vimentin positivity.

### Statistical analysis

Statistical analyses were performed with GraphPad Prism software 8.0. and R 3.6.3. Growth rates were analyzed by response feature analysis through fitting the growth data for each mouse individually to an exponential growth model, obtaining estimates of the rate parameters via nonlinear least squares. The growth rather were then compared using a one-way ANOVA with Tukey’s honest significant differences used to correct for multiple comparisons. Survival data was analyzed using the Kaplan-Meier method. Pathologic data was qualitatively characterized and analyzed using Chi-squared tests with a Bonferroni correction for multiple hypothesis testing.

## Results

### Prostate specific deletion of Trp53 and Pten leads to fast-growing, lethal, sarcomatoid tumors

To non-invasively monitor tumor burden, bioluminescence imaging (BLI) was performed on mice with Probasin-Cre mediated deletion of *Pten* and *Trp53*, coupled with the activation of a ROSA-LSL luciferase reporter. In this study, all mice had homozygous floxed *Pten* alleles, with cohorts having wildtype *Trp53* (WT), heterozygous floxed/wildtype *Trp53* (HET), or homozygous floxed *Trp53* alleles (DKO). The longitudinal BLI imaging (**Fig 1 A, B**) demonstrated that tumors grew faster in the DKO mice than those in the HET and WT group (**Fig S1**). After an initial rapid growth phase in all groups from 5-10 weeks, which reflects pubertal activation of Cre and growth of the prostate, WT mice showed little growth out to 50 weeks consistent with our previous studies (9). The DKO mice showed rapid tumor growth around 19 weeks of age that resulted in formation of readily palpable primary tumors and a median survival of 26.5 weeks (**Fig 1B, C**). In comparison to the DKO group, the HET group demonstrated slower disease progression with tumor growth not increasing until mice were around 29 weeks of age (**Fig 1B**). Moreover, the median survival for heterozygous *Trp53* null mice was closer to that of WT (41 and 50 weeks, respectively) (**Fig 1C**).

**Figure 1.**
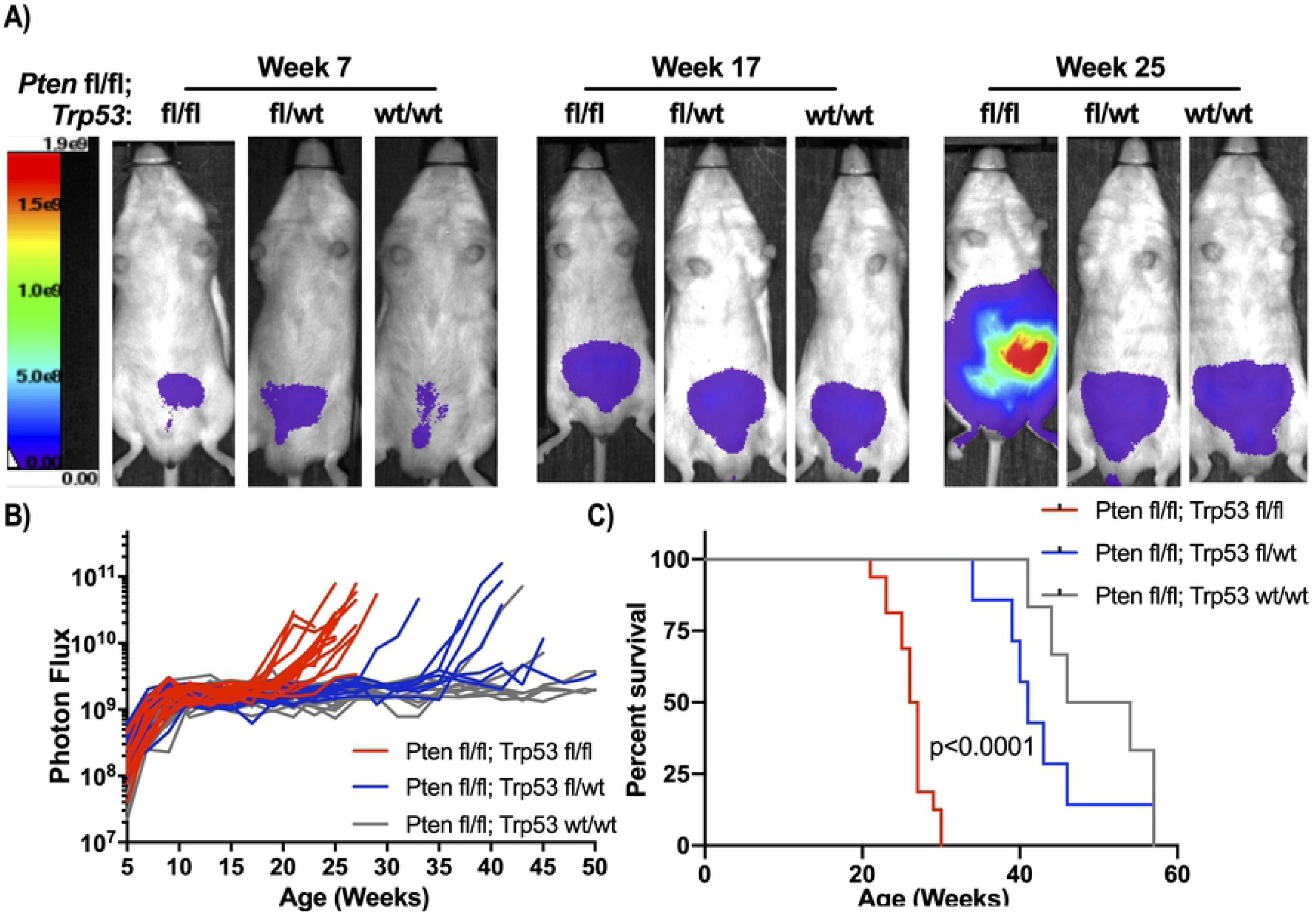
(A) Serial bioluminescence imaging (BLI) of mice with loss of *Pten* and wildtype *Trp53* (WT;wt/wt), heterozygous (HET;wt/fl), or homozygous (DKO;fl/fl) deletion of *Trp53*. (B) BLI signal development over time of each of the groups, BLI signal grew faster in the DKO than WT or HET mice (p<0.01). (C) Kaplan-Meier curves showing DKO mice have decreased survival compared to WT and HET mice (p<0.0001) (n=16 for DKO; n=7 for HET; n=6 for WT).

We performed histopathological analysis of prostates collected from DKO, HET, and WT mice euthanized at 25 weeks of age, as well as HET and WT mice at ∼40 weeks (**Fig 2A**). At 25 weeks, 13/13 mice from the DKO group demonstrated sarcomatoid tumors as confirmed by negative staining with a pan-cytokeratin antibody, with most of the HET and WT showing high-grade mouse prostate intra-epithelial neoplasia (HG mPIN) features. For the WT group, there was no difference in this distribution at 40 weeks; however, for the HET group, 6/6 tumors evaluated were sarcomatoid (**Fig 2B**). Based on the histological characterization, we determined if loss of Trp53 lead to elevated EMT-like features in the tumors. For this we stained tissue for the EMT markers Zeb1 and vimentin. For tumors to be scored positive for EMT we required that regions of overlapping positivity of both markers to be present. From this analysis, we observed an increase in EMT-positive tissue in the DKO group (at 25 weeks) and the HET group (at both 25 and 40 weeks) as compared to the WT group (**Fig 2C**).

**Figure 2.**
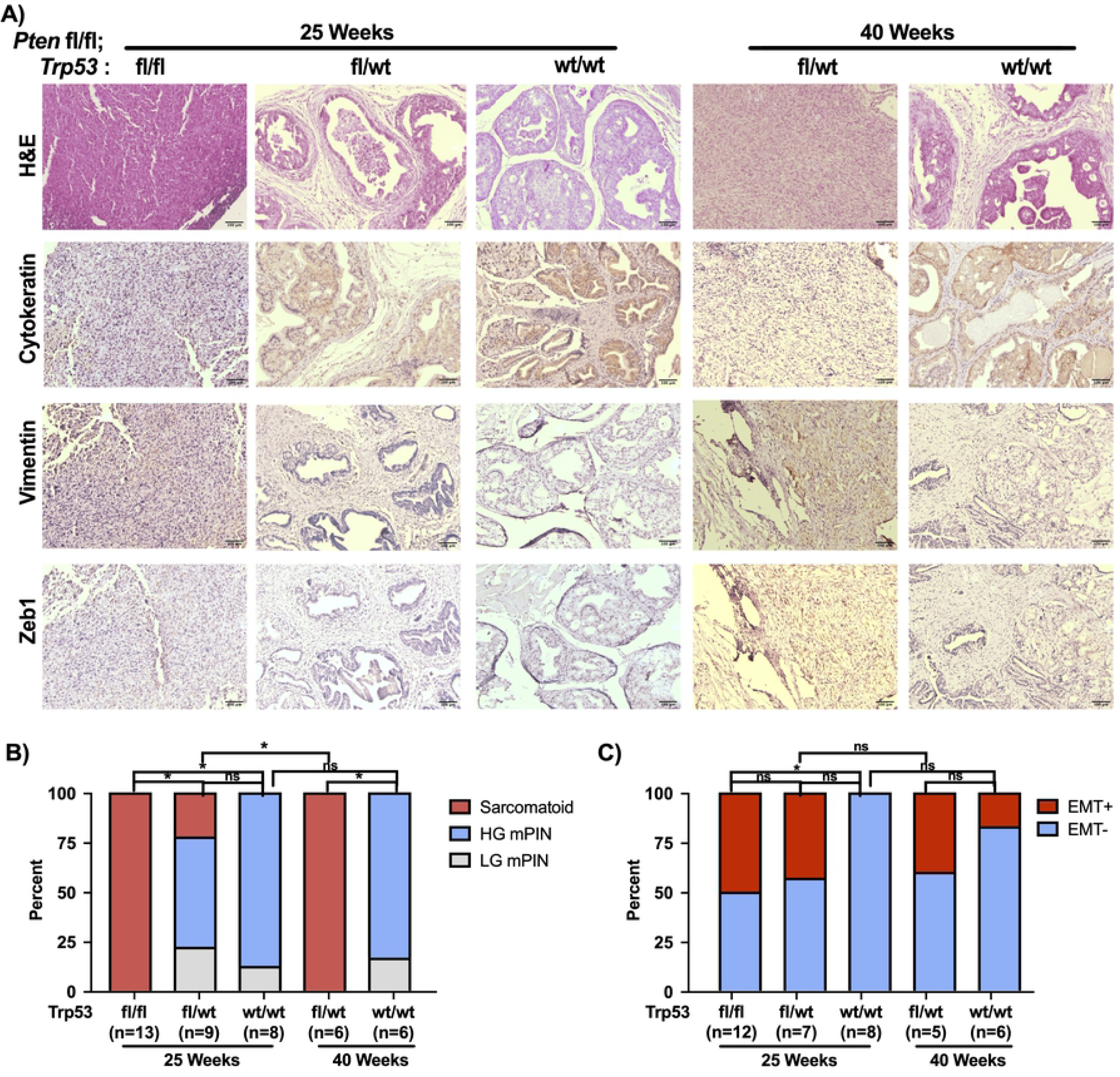
(A) Representative sections of tissue form DKO mice, as well as both HET and WT at 25 and ∼40 weeks. Disease stage was determined by pathologist review of H&E sections, with sarcomatoid being confirmed with a negative pan-cytokeratin stain. (B) Histopathology analysis from H&E and cyokeratin staining in (A), (*p>0.05; Chi-Square with Bonferroni correction). (C) Analysis of Zeb1 and Vimentin staining from (A), with EMT positive (EMT+) tissue having areas of overlapping Zeb1 and Vimentin staining (*p>0.05; Chi-Square with Bonferroni correction).

### Pten/Trp53 double knockout mice develop locally invasive disease but not lymphatic or distant metastasis

Due to the aggressive nature of the tumors from the DKO group and the prevalence of *PTEN* and *TP53* mutations in metastatic disease, we determined if there was metastatic disease in these mice. We evaluated residual tumor burden by BLI post-necropsy, after removal of the primary prostate tumor for 10 out of 16 mice (**Fig 3A, D, F**). This was necessary because the intense signal from the primary tumor could obscure weaker signals emanating from metastatic sites. We observed intraperitoneal spread in 8 out of 10 mice. We then histologically evaluated BLI signal-positive tissue to determine if there was metastatic disease (**Fig 3B, C, E, G**). Upon histological evaluation of a mouse that appeared to have lumbar lymph node involvement (**Fig 3A**), we found that instead of intralymphatic growth, the tumor invaded into the local perilymphatic fat (**Fig 3B, C**). DKO mice (8/10) were found to have visceral BLI signal upon post-necropsy imaging (**Fig 3D**), but this seems more likely to be peritoneal seeding and local invasion than hematogenously-disseminated disease (**Fig 3E**). There was also evidence of aggressive, invasive disease with BLI-positive tumor invading into pelvic sidewall musculature (**Fig 3F, G**).

**Figure 3.**
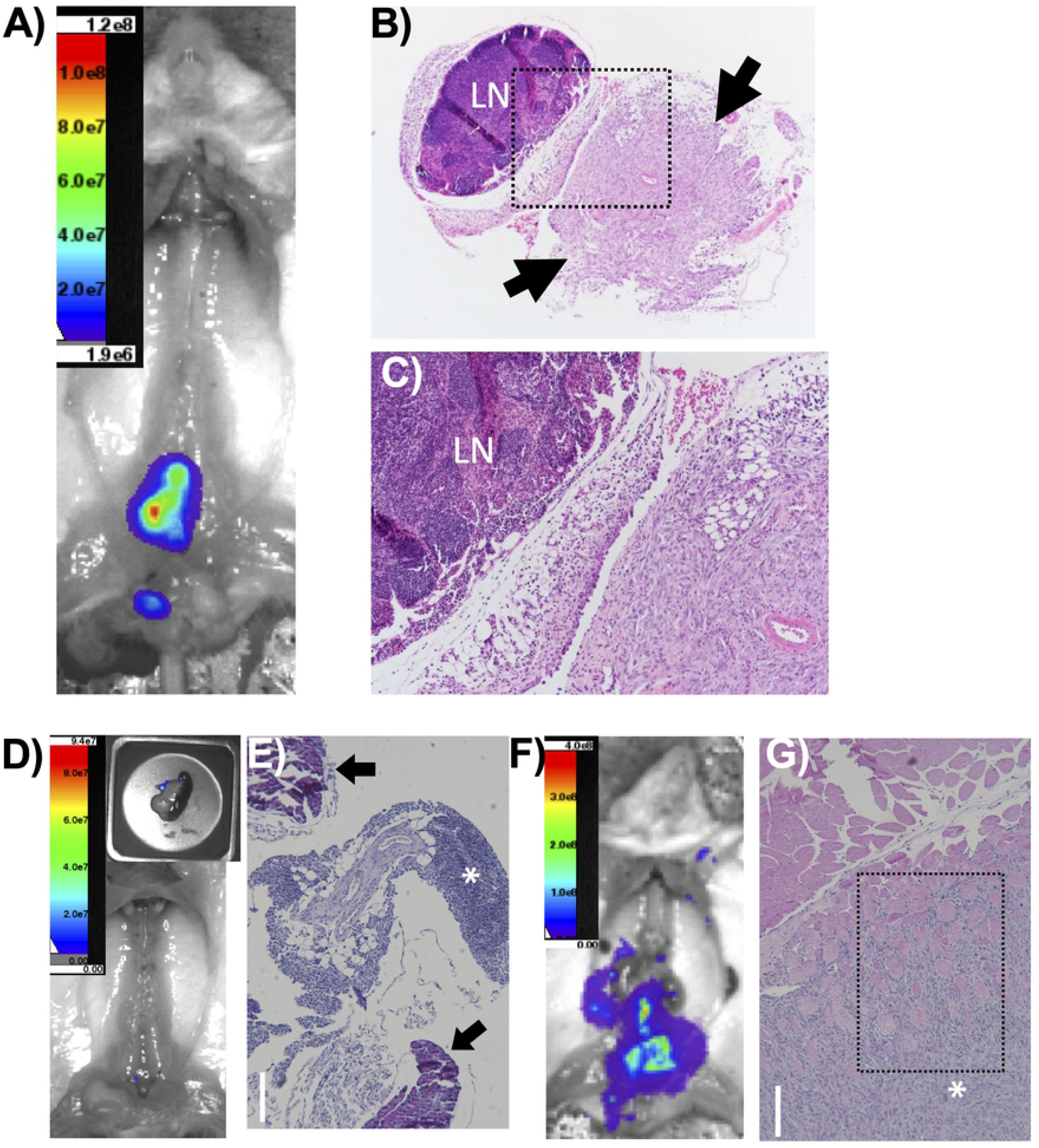
(A) Post-necropsy imaging in a DKO mouse. In situ, lymph nodes are BLI positive. (B-C) H&E stain showing a pelvic lymph node (LN) dissected at necropsy from the mouse in (A). The arrows represent areas of adjacent, locally invasive tumor. The area outlined by the black box in (B) is magnified in (C). (D) Post-necropsy imaging in a DKO mouse, with pancreas imaged separately (inset), with a nodule of tumor cells on the pancreas. (E) H&E stain showing the nodule dissected from the BLI-positive pancreatic lesion in (D). The tumor is indicated by the asterisk, and normal pancreatic tissue is marked with black arrows. The tumor is locally invasive into the tissue around the pancreas but not into the pancreatic tissue. (F) Post-necropsy imaging in a DKO mouse. The area near pelvic lymph nodes is BLI positive. (G) H&E stain of an area of BLI positivity in the mouse in (F). There is no lymphoid tissue, and the tumor (asterisk) is locally invasive into the adjacent muscle (black box).

### Pten/Trp53 double knockout results in loss of androgen receptor expression and castrate resistant tumors

While the DKO tumors were not metastatic, we wished to investigate the androgen sensitivity of this model. To assess this, we generated a cohort of five DKO mice which were surgically castrated at 10 weeks of age. After castration, the BLI signal noticeably decreased between 10 and 15 weeks of age (**Fig 4A**). Eight weeks after castration, the tumor burden started to increase in all mice, demonstrating castrate-resistant prostate cancer. Histological analysis from these tumors revealed that, like the intact DKO mice, they were all sarcomatoid (**Fig S2A, B**). Due to the rapid and uniform onset of castrate-resistant growth, we determined if the tumors from intact mice demonstrated alterations in AR expression (**Fig 4B, C**). From this analysis, we found that ∼75% of DKO mice lacked AR protein expression at 25 weeks, but all of the HET and WT mice retained AR expression at this timepoint (**Fig 4C**). At 40 weeks, ∼50% HET mice showed loss of AR expression. Consistent with the observation that tumors from the castrate DKO mice became androgen resistant, they also lacked AR expression (**Fig S2C**).

**Figure 4.**
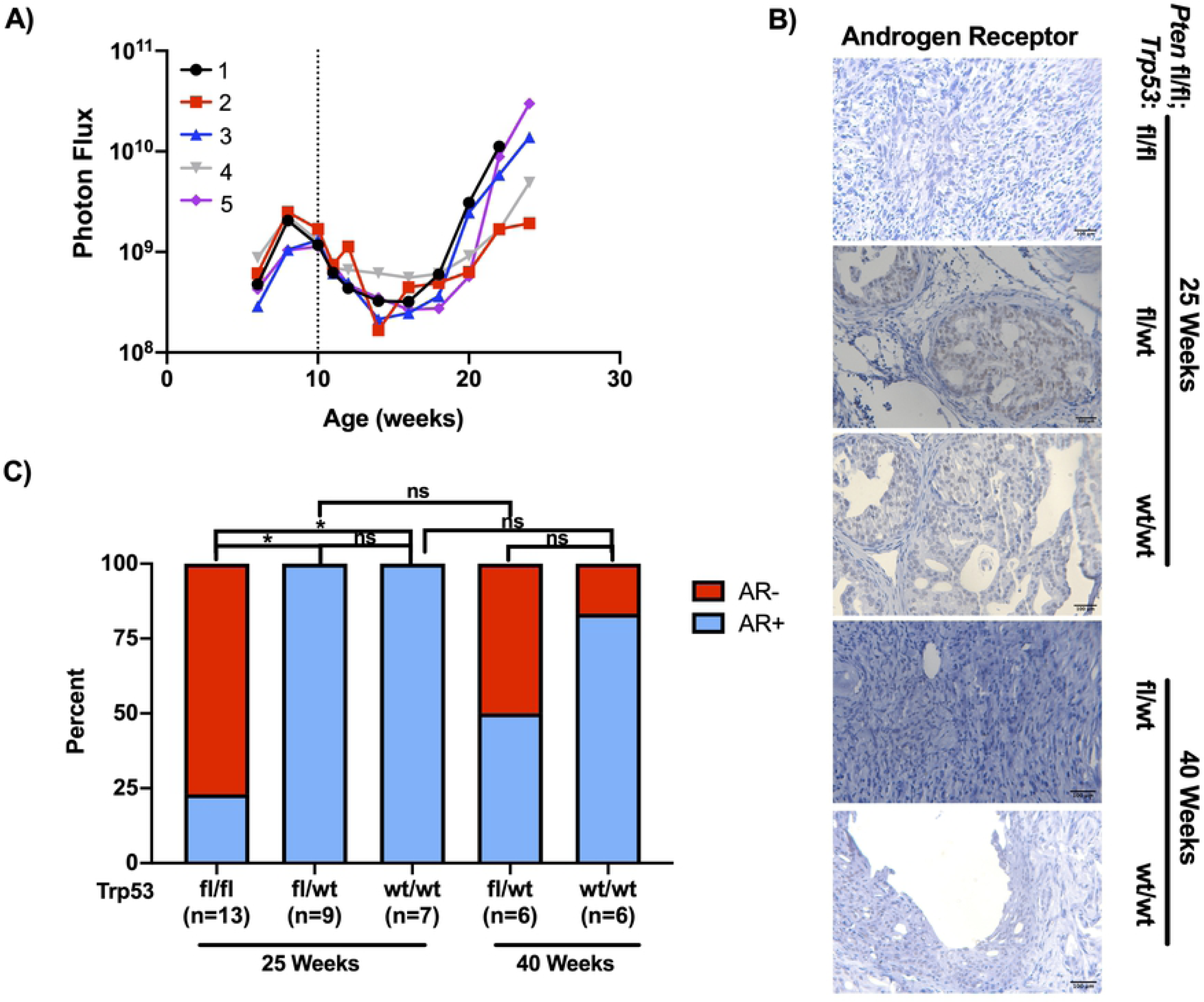
(A) BLI signal of a separate cohort of DKO mice that were castrated at 10 weeks of age (dotted line). Signal shows significant decrease immediately post-castration but over time recovers to levels of non-castrated counterparts. (B) Immunostaining of DKO, HET, and WT tissue for Androgen Receptor. (C). Analysis of (B) (*p>0.05; Chi-Square with Bonferroni correction).

## Discussion

Here we describe a Pb-Cre mediated *Pten*;*Trp53* double knockout mouse model of prostate cancer with a bioluminescent reporter for non-invasive monitoring of tumor growth. Our model showed accelerated tumor growth and mortality of the DKO mice as compared to HET and WT mice. While at approximately 25 weeks of age the HET mice were more like the WT knockout mice, analysis of tissue collected at necropsy demonstrated that eventually the HET mice had progressed and became similar to the double knockout mice. Necropsy tissue from both the DKO and HET mice showed that tumors had sarcomatoid pathology. This is a rare, undifferentiated and mesenchymal prostate pathology (<1% of prostate cancer tumors) that is considered to be highly aggressive (19-21). The loss of p53 function may be associated with the development of sarcomatoid tumors in patients. Case studies have reported strong nuclear accumulation of p53 in the sarcomatoid regions of the tumor, consistent with oncogenic dominant-negative or gain-of-function mutations in p53 (22), whereas adenocarcinoma typically lacks strong p53 staining (23, 24). In addition to the accumulation of p53, it has been demonstrated that the sarcomatoid portion of a prostate tumor lacks AR staining (25).

Sarcomatoid prostate cancer has been previously observed in *Trp53*;*Pten* prostate KO mice. In a report from Martin et. al. the authors noted that the progression of prostate tumors from Pb-Cre mediated knockout of *Trp53*;*Pten* resulted in a complete penetrance of spindle cell carcinoma or sarcomatoid pathology for mice between 26-30 weeks (11). They also noted that the regions containing spindle cell carcinoma had low or undetectable AR expression (11). Our results are consistent with these findings, as pathological analysis of necropsy tissue from our DKO mice displayed sarcomatoid features with most tumors lacking AR staining. We also demonstrated that terminal tissue from the HET mice were similar to DKO mice, as all tumors presented as sarcomatoid and 50% did not stain for AR. Previous studies with *Trp53*;*Pten* prostate KO mice did not report the androgen sensitivity of the autochthonous tumors. Utilizing BLI, we were able to determine that the DKO mice were initially responsive to castration but developed castrate-resistant disease 5 weeks later. The age for the onset of tumor growth in castrated DKO mice is consistent with the age in which the Martin study started to observe sarcomatoid pathology (11). Given the reduced AR protein expression in sarcomatoid cancers and that the castrate DKO mice all had sarcomatoid pathology, it is likely that onset of castrate-resistant tumor growth was due to the progression of the prostate cancer to a sarcomatoid pathology. While the development of sarcomatoid pathology under castrate levels of androgen is interesting, it is still unknown what events lead to the formation of this disease state. Understanding the progression to an AR-negative or -indifferent state may inform how prostate cancer bypasses AR signaling without developing a neuroendocrine (NE) phenotype, as the frequency of AR negative, non-NE prostate tumors is increasing (26).

Another interesting aspect of this model is the connection between the sarcomatoid pathology and EMT. In both the Martin study and case reports from patients, the sarcomatoid regions stained strongly for vimentin expression (11, 23). Vimentin, along with Zeb1, are widely regarded as markers of EMT and have been implicated in the development of metastatic disease in many cancers, including prostate (27). Our results show increased expression of both proteins in DKO tumors, and indeed we have shown that the expression of these proteins occurs within the same cell. Interestingly EMT-like characteristics may result from the reduction of AR signaling in the sarcomatoid tumors. It has been demonstrated that androgen deprivation leads to elevated expression of EMT markers in both normal and cancerous prostate tissue (28). Moreover, Zeb1 and AR may regulate each other through mutually antagonistic processes (28).

Despite the presence of EMT markers in the sarcomatoid tumors, neither in the earlier reports nor in our study were distant metastases detected (10, 11). Even with the addition of a bioluminescent reporter, we were able to detect only bioluminescent-positive tumor deposits that represented local invasion into peri-lymphatic fat as well as peritoneal seeding with tumor found on the pancreas. However, we cannot rule out the possibility that we did not detect micrometastatic disease in some organs which lacked sufficient BLI signal for detection because we did not exhaustively section metastatic end organs without BLI signal. In spite of the inability of BLI to detect metastatic disease in the model reported here, we believe there is utility in non-invasive, bioluminescent monitoring in the detection of distant, and potentially very small volume, metastatic disease. Moreover, the results from our study underscores the importance of validating BLI signal with histopathology.

For distant metastases to occur in the context of *Trp53*;*Pten* prostate-specific deletion, additional genes may need to be disrupted, such as *Rb1* or *Smad4* (12, 29, 30). However, expression of EMT markers Zeb1 and vimentin suggests metastatic potential in this model, and certainly correlates with more aggressive disease including castrate resistance.There are at least two aspects of this model which may contribute to the lack of observed metastatic disease. First, because *Trp53*;*Pten* are deleted in the entire prostatic epithelium, the subsequent tumors in this model are rapidly fatal, likely from mass effect and urinary obstruction, which limits the time for progression of disease. To circumvent these problems, somatic deletion of Pten and Trp53 may provide a new direction. Indeed, a virally-introduced, Cre-mediated deletion of *Pten* and *Trp53* in mouse prostates has reportedly produced reliable metastasis (30). Second, the host strain of the model may influence metastatic potential (6). It has been shown that normal prostate tissue from different strains have alterations in expression of genes such as MMP7 and prostate specific stem cell antigen (31). Moreover, BALB/c mice have loss of p16^Ink4a^, which could alter metastatic potential by dysregulation of the Rb pathway (32). Evaluating the loss of *Trp53*;*Pten* function across a range of host strains may elucidate this possibility and lead to the identification of metastasis modifier genes.

## Acknowledgements

We would like to thank Sophia Peterson for assistance with histopathology and Dr. Jennifer Richer for generously providing the Zeb1 antibody.

## Authorship

Conceptualization: NLB, RS, MDH; Data Acquisition: CY, DLM, NB, MV, JAB, MBC; Data Analysis: CY, DLM, PB, MBC, MDH; Writing (initial draft): CY, DLM, MDH; Writing (review/revisions): all authors.

## Supplemental Figure Legends

**Fig S1.**
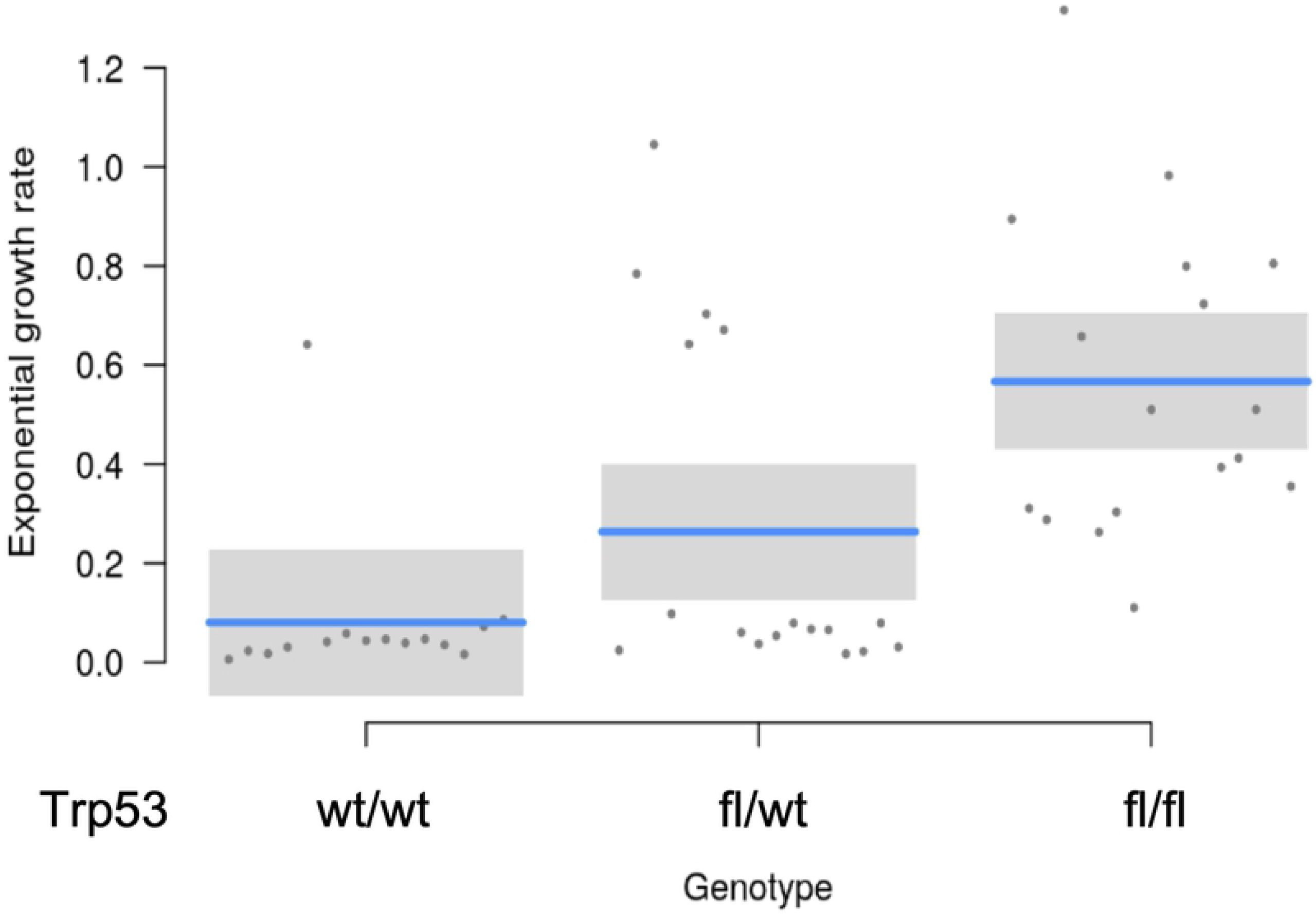
Double knockout mice have accelerated tumor growth. Graph of the exponential growth rate for each mouse from the Pten fl/fl; Trp53: wt/wt, fl/wt, and fl/fl groups. Blue line is the mean with grey boxes outlining the 95% confidence intervals.

**Fig S2.**
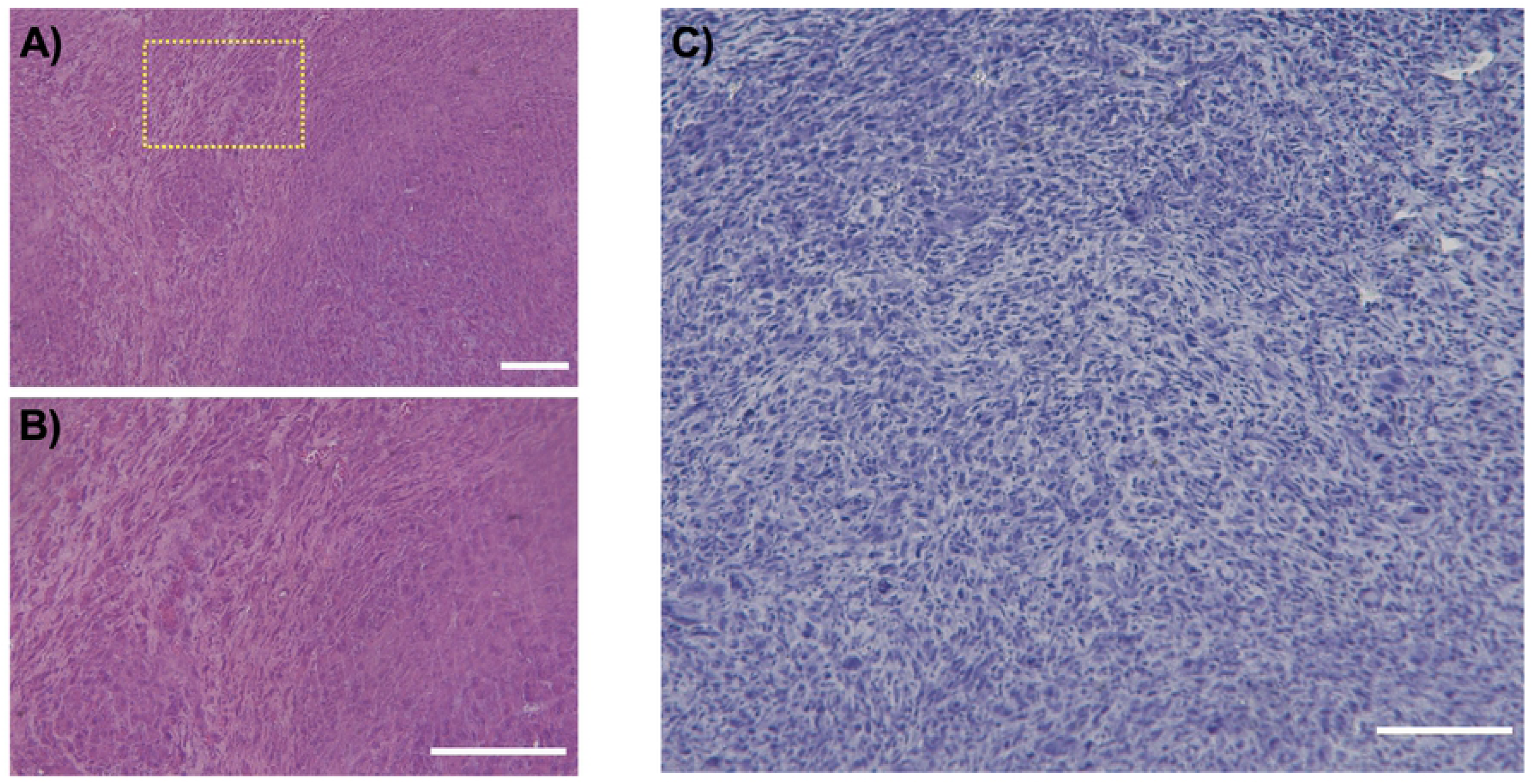
Histopathology of castrate DKO mice. (A, B) H&E stain showing sarcomatoid carcinoma, similar to the non-castrated DKO mice. Yellow box in (A) is enlarged in (B) to show sarcomatoid architecture. (C) Immunostaining for AR. Scale bar = 150 µm

